# The effects of fox movement and landscape heterogeneity on the spread of sarcoptic mange in urban settings

**DOI:** 10.64898/2026.06.24.734291

**Authors:** Nikol Dimitrov, Tiziana A. Gelmi-Candusso, Martin Krkošek, Marie-Josée Fortin

## Abstract

**Context:** The movement of vertebrate hosts across urbanized landscapes can play a key role in the transmission of direct-contact diseases. Understanding how wildlife hosts move in urban landscapes, and how transmission is affected by their landscape-constrained and disease-altered movements, is imperative for better predicting the spread of disease.

**Objective:** We assess how the movement of red foxes (*Vulpes vulpes*) according to landcover type, and their infection status, affect the spread of mange (caused by *Sarcoptes scabiei*) in an urbanized landscape.

**Methods:** We developed a mange transmission model (MTM) using an agent-based model to compare two movement behaviours of foxes in Scarborough (Ontario, Canada): random and landcover-based. We further assessed the effects of movement on disease transmission by considering the fox’s infection status and comparing a range of movement probability scenarios. We quantified the number of effective contact events and the effective reproduction number (*R*_e_) according to each scenario.

**Results:** We found that both landcover-dependent movement and infection status influenced the spread of mange within fox populations. The number of effective contact events and effective reproduction number *R*_e_ was greatest when landscape heterogeneity was included in the model and foxes moved through paths of least resistance to movement, and when susceptible and infected foxes had an equal probability of leaving a fragmented habitat patch.

**Conclusions:** Our findings suggest that mange spread may be accelerated along movement corridors in fragmented, heterogenous landscapes. As urban areas expand and remnant habitat within these is further lost and animals are relegated to fewer movement pathways, disease transmission may increase.

## 1 Introduction

In recent years, urban areas across North America and Europe have seen an increase in infectious diseases such as Lyme disease, chronic wasting disease, and rabies (Mather et al. 1996; Williams et al. 2002; Bradley & Altizer 2007; Beran 2017; Guo et al. 2022; Dubey et al. 2023). Host species for some of these diseases, such as red foxes (*Vulpes vulpes*), coyotes (*Canis latrans*), and raccoons (*Procyon lotor*), have adapted particularly well to living in urban cores, leading to an increased risk of spread within wildlife populations and spillover infections to other hosts (Guillot et al. 2024). Red foxes, in particular, are distributed throughout over 110 cities in North America, Europe, the UK, and Australia (Soulsbury et al. 2010; Lombardi et al. 2017). These increases have been mainly attributed to their ability to utilize urban features, and anthropogenic food waste, for their sheltering and dietary needs (Contesse et al. 2004). The increase in urban fox densities has led to higher transmission of diseases including more frequent disease outbreaks (Plumer et al. 2014). In addition, in urban settings, there is evidence of fox habitat overlap with coyotes, raccoons, and gray squirrels (*Sciurus carolinensis*), which can also carry compatible diseases, such as mange (Mueller et al. 2018).

Sarcoptic mange is a highly infectious disease caused by the parasitic mite *Sarcoptes scabiei* (Soulsbury et al. 2007). *S. scabiei* is a host generalist with a microparasitic lifestyle, that has impacted over 140 different species of both wild and domestic mammals worldwide (Scott et al. 2020; Escobar et al. 2022; Guillot et al. 2024) Infected mammals develop skin lesions, hyperkeratosis, and erythema and may succumb to secondary infections (Scott et al. 2020). A high prevalence of the parasite has led to severe reductions in mammal populations, including an over 90% reduction in a Japanese population of wild raccoon dogs (*Nyctereutes procyonoides*), declines in litter birth rates in a population of infected arctic foxes (*Vulpes lagopus*), and a 50% reduction in a California population of kit foxes (*Vulpes macrotis*) (Foley et al. 2023; Matsuyama et al. 2024; Wallén et al. 2024). Mange is transmitted through direct contact with an infected host or indirect contact through a contaminated environment (Arlian & Morgan 2017). Though environmental transmission from den sharing has been a noted hypothesis, the primary mode of transmission observed in foxes has been direct contact between infected individuals (Montecino-Latorre et al. 2019; Scott et al. 2020). Disentangling the respective impacts of these modes of transmission remains a challenge for den-using mammals (Martin et al. 2018).

Periodic outbreaks of mange have led to fox population declines in Europe, with a slow recovery phase (Baker et al. 2000; Soulsbury et al. 2007; Wallén et al. 2024). Such epizootics have resulted in a disruption of community dynamics where prey species of foxes increased in abundance (Lindström et al. 1994). Transmission between species can also occur, amplifying the problem across the urban wildlife community (Kołodziej-Sobocińska et al. 2014). Recent findings on the expansion of mange to a variety of host species, stress the urgency of studying mange as an emerging wildlife disease (Guillot et al. 2023). Consequently, understanding how mange may spread across fox individuals in urban settings is becoming increasingly important from an ecological perspective.

Variation in landcover types within cities, ranging from natural green areas to highly anthropogenic areas such as commercial and residential areas, influences animal movement (Cadenasso et al. 2007). Foxes are less likely to utilize heavily impacted human regions (Duduś et al. 2014), and more likely to move through less human-impacted landcover types, which can sometimes create movement corridors between urban green fragments (Aziz & Rasidi 2014; Gallo et al. 2017). Foxes have been noted to utilize smaller, fragmented residential parks in the city of Brighton, UK, suggesting these green areas play a critical role in habitat selection of urban foxes (Tolhurst et al. 2020). Increased animal movement between habitat patches generally leads to enhanced pathogen transmission (Hess 1996), therefore landscape heterogeneity and resulting habitat connectivity, which influence animal movement, may alter disease dynamics. For example, landscape connectivity can increase the risk of chronic wasting disease in deer (Norbert et al. 2016), stepping-stones between habitat patches can enhance the spread of ticks carrying Lyme disease (Watts et al. 2018, Heylen et al. 2019), landscape permeability and movement directionality can increase the spread of feline immunodeficiency virus in bobcats (*Lynx rufus*) (Fountain-Jones et al. 2017), and connectivity can act synergistically with proportion of forest cover in defining contact rates for urban raccoons (Tardy et al. 2018). Therefore, understanding how foxes interact with the urban landscape from a connectivity perspective could prove particularly useful for more accurate quantification of contact events and mange dynamics. Connectivity assessments are often based on a cost surface function and are particularly useful for the effective modelling of movement behaviours in fragmented heterogeneous landscapes (Rayfield et al. 2010; Etherington 2016). These connectivity assessments are derived from resistance to movement values, which depend on the ability of species to move through different landcover types (Doerr et al. 2011). Incorporating such movement costs into models of fox movement in urban landscapes could help identify areas in the city where mange infections are more likely to occur and how this affects transmission dynamics.

In addition to landscape heterogeneity, mange infection itself may also alter the movement behaviours of its hosts. For example, mange-infected wombats are more likely to spend time outside of their burrows and less time moving around their habitat (Simpson et al. 2016). Similarly, mange-infected foxes may behave abnormally. A comparison between healthy and mange-infected foxes found that mange-infected foxes have more abnormal patterns of movement and tend to move through normally less used landscape types (Overskaug 1994). Yet, previous studies that have modelled mange dynamics, have not considered how variable movement abilities of susceptible versus infected foxes may influence mange spread (Lunelli 2010; Devenish-Nelson et al. 2014). These differing movement abilities may lead to unequal levels of contact rates within habitat patches and along movement paths (Foley et al. 2023). Including both differing movement abilities between infected and susceptible individuals and resistance-to-movement values informed by behavioural patterns of urban foxes, can provide a more accurate representation of mange infection dynamics in cities.

Mange dynamics have mostly been modelled using compartmental models, in particular, SEI (Susceptible-Exposed-Infected) models, as infected individuals who recover do not build up immunity to secondary infection (Kermack & McKendrick 1927; Lunelli 2010; Devenish-Nelson et al. 2014). While compartmental models can reveal important population-level phenomena they also tend to ignore spatial properties and assume a homogenous mixing of susceptible and infected individuals (Smith & Moore 2004). Unlike these models, agent-based models (ABM) incorporate spatial variation in transmission, which allows for complex population-level phenomena to arise from individual behaviours (Judson 1994). ABMs have been used extensively for modelling realistic dispersal behaviours of wildlife species and have also been employed for dynamic modelling of disease spread (Perez & Dragicevic 2009; Tang & Bennett 2010). ABMs are therefore ideal tools for incorporating realistic host movement behaviours considering connectivity and disease-dependent movement patterns into models of mange.

In this study, we assess how fox movement influences mange transmission in the urban context, by incorporating landcover structure and infection-dependent movement ability into an ABM where fox movement occurs between two habitat patches. We use this model to assess:(1) The influence of landscape heterogeneity on the effective reproduction number (*R_e_*) and the effective contact events between hosts. (2) The influence of movement ability of susceptible vs. infected foxes on the transmission of mange. We hypothesize that landscape heterogeneity will exacerbate spread, leading to higher *Re* and effective contact events among foxes, particularly along movement corridors between habitat patches. In addition, we hypothesize that increased movement of infected foxes outside of green areas will increase transmission across the landscape to susceptible hosts. Addressing these gaps in knowledge of the spread dynamics of mange and identifying urban areas where disease transmission is more likely to occur, will provide useful information to better target disease control efforts in urban environments.

## 2 Methods

### 2.1 Study area

Our model was applied to a 1×1km urban portion of the Scarborough area in Toronto (Canada), using a cell resolution of 10×10m (Figure 1). Due to the high computational demands of our model, we focused on a smaller study area, which allowed us to capture urban spatial heterogeneity at a finer resolution. The study area was selected due to its high level of landscape heterogeneity, consisting of eight landcover types which included highly urbanized areas such as commercial and residential locations, natural areas such as parks, and linear features such as roads. Within this area, our model used two parks: Warden Woods Park and Runnymede Lands Park. This area was selected as foxes are expected to utilize parks has habitat patches and move within and between them. Landcover information was compiled using ArcGIS and data from the TRCA (ESRI 2011; Toronto and Region Conservation Authority, 2017).

**Figure 1.**
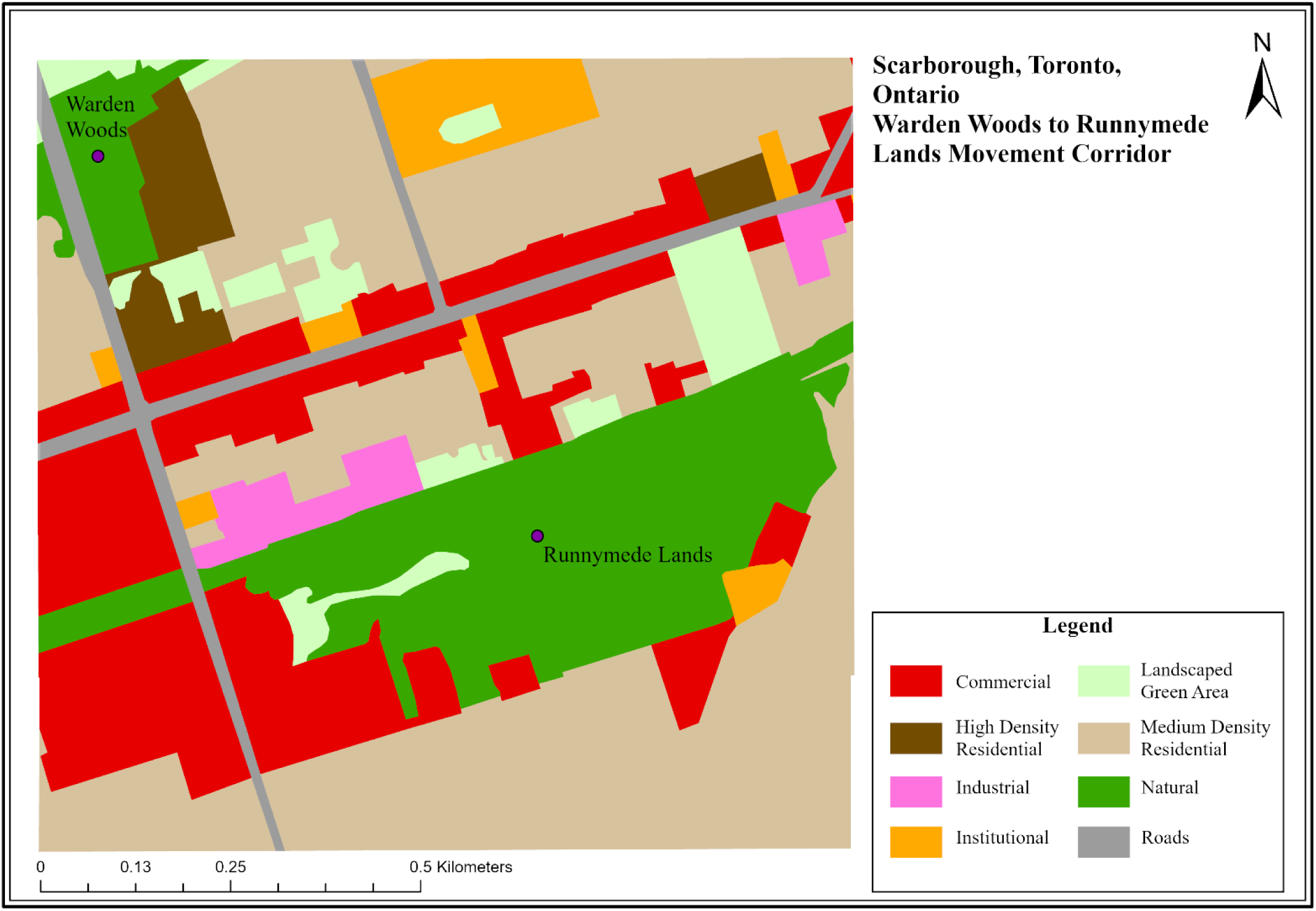
Warden Woods to Runnymede Lands movement corridor in Scarborough, City of Toronto, Ontario, with landcover types following the Toronto Region Conservation Authority Landuse NaturalCover map from 2017 (TRCA, 2017)

### 2.2 Model Description

We developed an agent-based model——using NetLogo 6.1 (NetLogo, 1999) and we called this model the Mange Transmission Model (hereafter MTM). The MTM is a discrete-time, spatially explicit model containing three main submodels: (1) a within-park movement procedure, (2) a between-park movement procedure, and (3) a disease procedure. The components of the three MTM submodels are described following the ODD (Overview, Design Concepts, Details) protocol (Appendix: ODD Protocol; Grimm et al. 2010). Foxes followed the SEI disease stages, with healthy foxes being classified as susceptible (S), foxes who have come in contact with diseased foxes but are not yet symptomatic classified as exposed (E), and diseased foxes classified as infected (I). Demographic factors of foxes, and seasonality, were not considered in our model. Using this model, we assessed (1) the effect of landscape heterogeneity on disease spread following the inclusion landcover resistance values (2) the effects of infection status on the spread following differential movement ability.

### 2.3 Movement Analysis

In our model analysing the effect of landscape heterogeneity on disease spread, we compared the effects on transmission following two movement behaviours within the study area for a population of 16 foxes: (1) random movement where fox movement is not affected by the landcover type (i.e., Mr), and (2) landcover-based movement (i.e., Mlb) where fox movement was facilitated or impeded according to the landcover types and their associated resistance values. The resistance values for the latter scenario were calculated across the landcover types based on four different factors that could impact the movement of urban foxes:human presence, temporal human activity, the degree of natural vegetation cover, and landscaping level, with the underlying assumption that foxes avoid human presence and will prefer sheltered areas to move across the landscape. These factors were assigned a numerical value, described in Table 1, based on camera trap observations we conducted in our study area (Supplementary Material: Table S3), and their arithmetic sum was used as a resistance value for each landcover type. These resistance values were further evaluated through a sensitivity analysis in James (2021) where relative differences played a larger role in determining least cost paths as opposed to absolute values, in accordance with previous studies (Rayfield et al. 2010, Albert et al. 2017). To add natural variability in resistance across the landscape, especially within the same landcover type, a random float value between 0 and 1 was added to each resistance value in the study area. We developed this novel approach to enable more natural movement of foxes across contiguous cells of the same landcover type and allow for the outflow of foxes from the least resistant areas, i.e., the forested patches.

**Table 1.** Resistance values of eight landcover types in Scarborough region that influence fox movement in urban settings (Total Resistance Value) based on how different components of the landscape (Human presence, Vegetation Cover, Landscaping Level [i.e. whether vegetation is actively managed (high)] and Temporal Human Activity [i.e. temporal usage patterns of landcover type by people]) within each landcover type can create resistance to animal movement. Numerical values in brackets represent the resistance value associated with the landscape factor affecting movement. These values are then summed up to result in the total resistance value for each landcover type. Sensitivity of resistance values was examined in James (2021).

| Landcover Type | Human Presence | Vegetation Cover | Landscaping Level | Temporal Human Activity | Total Resistance Value |
| --- | --- | --- | --- | --- | --- |
| Commercial | High (25) | Open (10) | High (25) | Daily (25) | 85 |
| High-Density Residential | High (25) | Low (25) | High (25) | Weekends (1) | 76 |
| Industrial | High (25) | Low (25) | High (25) | Weekdays (10) | 85 |
| Institutional | Medium (10) | Open (10) | High (25) | Weekdays (10) | 55 |
| Landscaped Green Area | Low (1) | Open (10) | High (25) | Weekends (1) | 37 |
| Road | High (25) | Heterogenous (1) | High (25) | Daily (25) | 76 |
| Medium-Density Residential | Medium (10) | Heterogeneous (1) | High (25) | Weekends (1) | 37 |
| Natural (including two parks and park edges) | Low (1) | Heterogenous<br>(1) | Low (1) | Weekends<br>(1) | 4 |

In our analysis of the impact of movement ability on disease spread, foxes were assigned a probability of leaving a park according to their infection status (i.e., whether they were susceptible, exposed, or infected). This parameter was then varied in the model, resulting in three scenarios with varying intensities: (1) an equal probability of leaving a park at a low intensity (i.e., eq-l), at a medium intensity (i.e., eq-m) and at a high intensity (i.e., eq-h), (2) a susceptible-biased probability of leaving a park at medium (i.e., sb-m) and low intensities (i.e., sb-l), and (3) an infected-biased probability of leaving a park at medium (i.e., ib-m) and low intensities (i.e., ib-l) (Table 2). These scenarios followed previous studies finding differences in movement abilities between mange-infected canids and susceptible individuals, where Mange-infected canids were more likely to utilize anthropogenic features and move greater distances (Murray et al. 2015), mange-infected canids in the acute stage, were less mobile following energetic losses related to their immune response (Nimmervoll et al. 2013).

**Table 2.** Scenarios tested for two movement behaviours: random movement and landcover-based movement: and the probability of leaving a park based on infection status: susceptible (S) (grouped with exposed E) and Infected (I). For the equal probability of leaving a park, 3 different intensities of the parameters were tested: low, medium, and high. For the infected-biased probability of leaving a park, the infected probability was kept high, with the susceptible (S) and exposed (E) probabilities tested at low and medium intensities. For the susceptible-biased probability of leaving a park, the susceptible probability was kept high, with the infected (I) and exposed (E) probabilities tested at low and medium intensities.

|  | Equal Probability of leaving park<br>(eq-) |  |  | Infected-Biased<br>Probability of leaving<br>park<br>(ib-) |  | Susceptible-Biased<br>Probability of leaving<br>park<br>(sb-) |  |
| --- | --- | --- | --- | --- | --- | --- | --- |
|  | Low | Medium | High | Low | Medium | Low | Medium |
| **Random Movement**<br>**(Mr)** | S & E – 0.25<br>I – 0.25<br>(Mr-eq-l) | S & E – 0.5<br>I – 0.5<br>(Mr-eq-m) | S & E – 0.75<br>I – 0.75<br>(Mr-eq-h) | S & E – 0.25<br>I – 0.75<br>(Mr-ib-l) | S & E – 0.5<br>I – 0.75<br>(Mr-ib-m) | S & E – 0.75<br>I – 0.25<br>(Mr-sb-l) | S & E – 0.75<br>I – 0.5<br>(Mr-sb-m) |
| **Landcover-based Movement**<br>**(Mlb)** | S & E – 0.25<br>I – 0.25<br>(Mlb-eq-l) | S & E – 0.5<br>I – 0.5<br>(Mlb-eq-m) | S & E – 0.75<br>I – 0.75<br>(Mlb-eq-h) | S & E – 0.25<br>I – 0.75<br>(Mlb-ib-l) | S & E – 0.5<br>I – 0.75<br>(Mlb-ib-m) | S & E – 0.75<br>I – 0.25<br>(Mlb-sb-l) | S & E – 0.75<br>I – 0.5<br>(Mlb-sb-m) |

### 2.4 Disease transmission metrics

Each movement scenario (Mr-eq-l, Mr-eq-m, Mr-eq-h; Mr-sb-l, Mr-sb-m; Mr-ib-l, Mr-ib-m; Mlb-eq-l, Mlb-eq-m, Mlb-eq-h; Mlb-sb-l, Mlb-sb-m; Mlb-ib-l, Mlb-ib-m) in the MTM was replicated 100 times for a simulation duration of 1500 days where the following metrics were computed: First, we assessed the effective reproduction number *R_e_* for the endemic phase of the disease spread—i.e., the number of foxes infected by an infected individual during the last 1000 days of the simulation, excluding the initial disease invasion stage (first 500 days of the simulation time). The endemicity threshold was determined by an observation of the simulation behaviour, where spread was likely to either be sustained or die off around the 500-day mark. The *R_e_*, rather than the *R_0_*, was computed as the *R_e_* tends to represent a more accurate measure of infection in a population where both susceptible and infected individuals exist at a given time, accounting for the existence of mange in an endemic state within fox populations (Nishiura & Chowell 2009; Foley et al. 2023). In contrast, the basic reproduction number, *R_0_,* is calculated at the beginning of disease invasion, when one individual is infected, and the rest are susceptible. The *R_e_* was calculated by keeping a count of all infected foxes and the number of foxes they infected, which returned to zero when the individual recovered and became susceptible again (Wadkin et al. 2024). This value was then averaged for the whole population for the last 1000 days of the simulation. The second metric computed was the number of effective contacts according to landcover type—i.e., the number of times successful transmission occurs between a susceptible and infected individual. We then compared: (1) the number of simulations where spread occurs and was sustained throughout the simulation time—to assess how many simulation runs, out of 100, mange spread persisted (2) mean *R_e_*—summarized for all simulation runs where spread was sustained, (3) the number effective of contacts (i.e., any event where successful transmission occurs between a susceptible and infected fox) summarized for all simulations where spread is sustained (4) the proportion of effective # of contacts that occur within a park—i.e., the number of effective contacts within the park and park edge landcover types divided by the total number of effective contacts, and finally (5) the proportion of the effective number of contacts that occur outside of a park—i.e., the number of effective contacts within the non-park landcover types divided by the total number of effective contacts.

### 2.5 Data Analysis

Output data were aggregated and analyzed using R Statistical Software (v4.2.1; R Core Team 2021) and the *dplyr* (v1.3.1, Wickham et al. 2023), *tidyr* (v1.3.0, Wickham et al. 2023), and *janitor* (v2.1.0; Firke 2021) packages. Boxplots were created using the *ggplot2* (v3.4.0; Wickham 2016) package, and the mixed model estimate plot was created using the *jtools* (v2.2.0; Long 2022) and *sjPlot* (v2.8.12; Lüdecke 2022) packages.

To assess the effects of two movement behaviours and the probabilities of leaving a patch according to infection status on the *R_e_*, we fit a general mixed effects model with a Poisson distribution using the *glmer* function from the *lme4* (v3.1-3; Bates et al. 2015) package. This was done at the individual fox level with movement behaviour, infection status scenarios, and their interaction being assessed as fixed effects. Replication (i.e., run number), individual fox ID, and the number of times each fox became reinfected were nested and assessed as random effects. The number of times each fox became reinfected was set as a random effect, as there is a lack of genotype and phenotype effects in the model and then nested within fox identity due to the possibility of spatial autocorrelation along movement routes in the landscape. We set the optimization parameter nAGQ to 0 to decrease the computational time needed for processing the resulting large dataset (∼122,000 data points). We then assessed the significance of our model with an analysis of deviance using the *Anova* function from the *car* (v3.1-1; Fox & Weisberg 2019) package.

To assess the effects of the two movement behaviours and the probability of leaving a patch on the number of effective contact events, we fit a population-level general mixed model using a Poisson distribution, with movement behaviour, infection status, and their interaction as fixed effects and run number as a random effect. We then also performed an analysis of deviance to assess the significance of the results.

## 3 Results

When the movement behaviour between parks was random (i.e., no resistance to movement), the spread was sustained at a range of 69-85 simulations out of 100 (Supplementary Material: Table S1). Within the random movement scenario, the greatest number of simulations where spread was sustained was observed when susceptible and infected foxes had an equal probability of leaving a park at a low intensity (Mr-eq-l), for a total of 85 out of 100 simulations. In comparison, for the landcover-based movement, the number of simulations where spread occurred ranged between 75-83 out of 100, with the greatest number of simulations resulting in sustained spread occurring for the susceptible-biased scenario with medium intensity (Mlb-sb-m), for a total of 83 out of 100 simulations(Supplementary Material: Table S1). The dynamics of the mange epidemic were similar across the simulation runs, with a rapid increase in infection prevalence at the beginning of the simulation, i.e., the invasion stage, which was then maintained through the simulation time with small oscillations, leading to a stable endemic stage. Susceptible and exposed foxes were maintained at small numbers through the simulation time (Figure 2).

**Figure 2.**
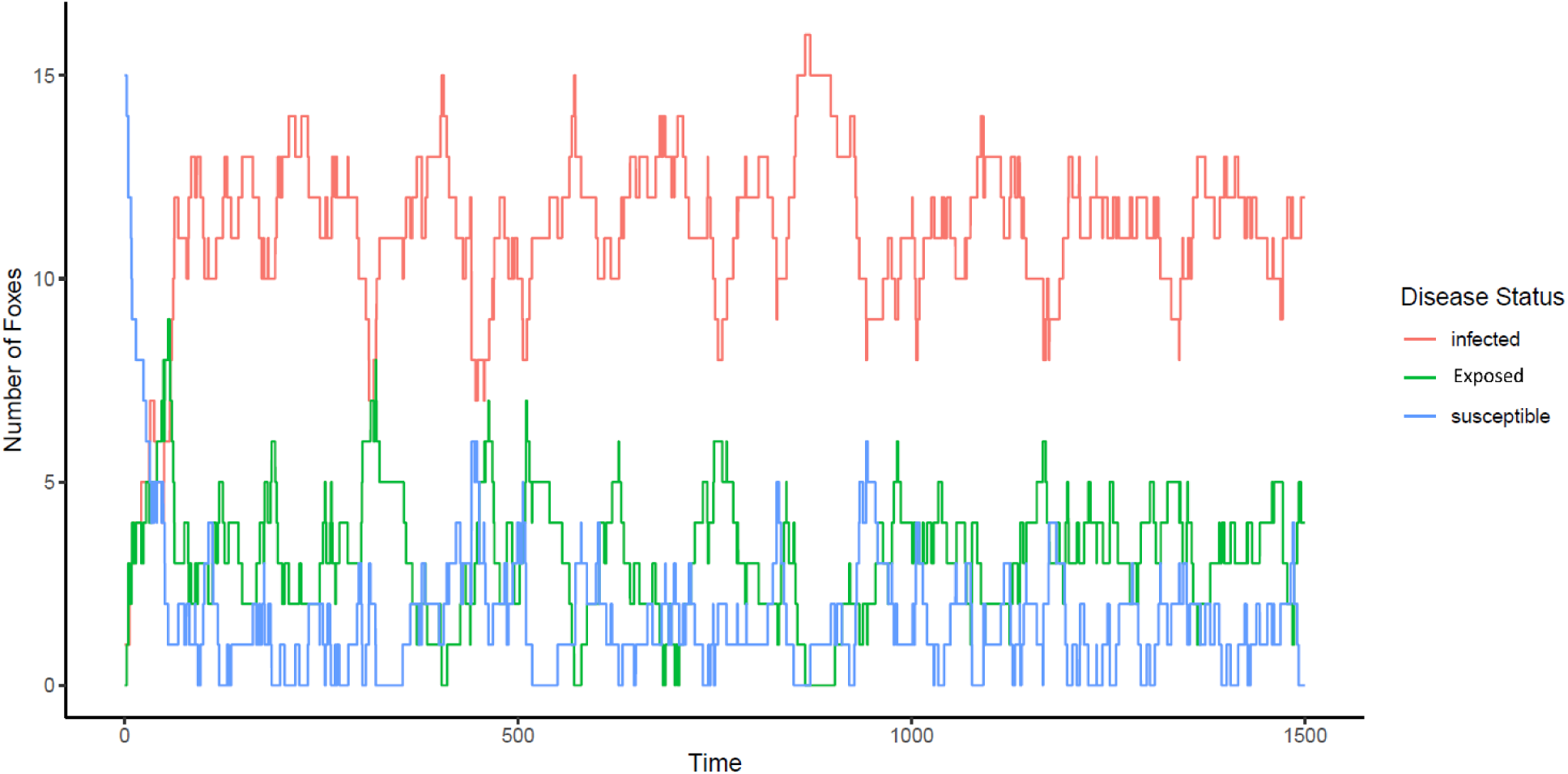
Number of foxes plotted versus simulation time for one simulation run according to infection status: infected (red), exposed (green), and susceptible (blue).

The mean *R_e_* computed for the landcover-based movement behaviour was 3.42 (± 0.61), compared to a mean *Re* of 2.12 (± 0.32) for when the movement behaviour was random (Supplementary Material: Table S1). Within both of the random and landcover-based movement scenarios, the mean *R_e_* was greatest when susceptible and infected foxes had an equal probability of leaving a park (Mr-eq; Mlb-eq) in comparison to the susceptible- and infected-biased probabilities (Mr-sb, Mr-ib; Mlb-sb, Mlb-ib), with the random scenario being an average 0.1 (± 0.2) greater, and the landcover-based scenario 0.4 (±0.14) greater. There was a small difference between the susceptible and infected biased scenarios in terms of mean *R_e_*, with an average difference of 0.01 for the random movement scenario, and 0.05 for the landcover-based movement scenario. However, the mean *Re* was greater when the parameters of leaving were more similar between susceptible and infected individuals (i.e., a smaller difference in the medium-intensity scenario between susceptible and infected foxes), rather than the low-intensity scenarios where there was a greater difference in the probability of leaving between susceptible and infected foxes (Figure 3(a); Supplementary Material: Table S1). The general mixed effects model showed a negative effect on the *R_e_* when movement was random in comparison to when movement was landcover-based, with landcover-based movement resulting in a greater *R_e_* regardless of the probabilities of leaving a park (Figure 3(a); Table 3). In addition, the infected-biased and susceptible-biased movement behaviours resulted in a lower *R_e_* in comparison to when movement between infected and susceptible foxes was equal. Results of the analysis of deviance for the model revealed a significant effect (*p* <2.2e-16) of movement behaviour and the probability of leaving a park according to infection status on the *R_e_* (Supplementary Material: Table S2).

**Figure 3.**
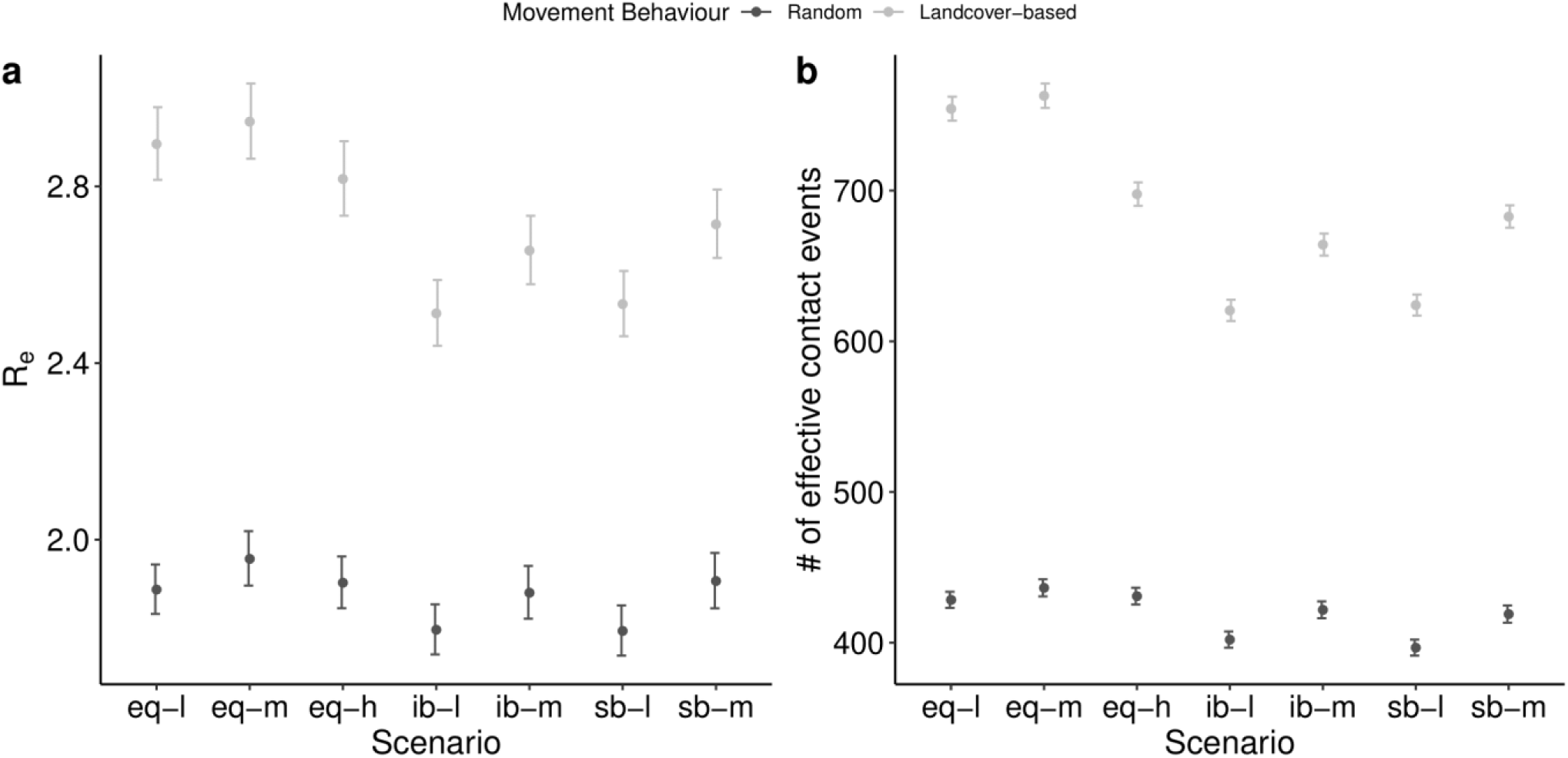
Grouped boxplot depicting **(a)** *Re* (summarized for the last 1000 days) and **(b)** number of effective contact events for the simulations where spread was sustained for the two movement behaviours: random and landcover-based on the general mixed model. Results are grouped according to the scenarios of leaving a park according to infection status: equal with low intensity (eq -l), equal with medium intensity (eq-m), equal with high intensity (eq-h), infected-biased with low intensity (ib-l), infected-biased with medium intensity (ib-m), susceptible-biased with low intensity (sb-l), and susceptible-biased with medium intensity (sb-m).

**Table 3.** Results for the *R_e_* general mixed model.

| *Predictors* | *Re* | *CI* |
| --- | --- | --- |
| Landcover based: equal-high | 2.82 | 2.74 – 2.90 |
| Landcover based: equal-low | 2.90 | 2.82 – 2.98 |
| Landcover based: equal-medium | 2.95 | 2.86 – 3.03 |
| Landcover based: infected biased-low | 2.51 | 2.44 – 2.59 |
| Landcover based: infected biased-medium | 2.66 | 2.58 – 2.73 |
| Landcover based: susceptible biased-low | 2.53 | 2.46 – 2.61 |
| Landcover based: susceptible biased-medium | 2.71 | 2.64 – 2.79 |
| Random: equal-high | 1.90 | 1.85 – 1.96 |
| Random: equal-low | 1.89 | 1.83 – 1.94 |
| Random: equal-medium | 1.96 | 1.90 – 2.02 |
| Random: infected biased-low | 1.80 | 1.74 – 1.85 |
| Random: infected biased-medium | 1.88 | 1.82 – 1.94 |
| Random: susceptible biased-low | 1.79 | 1.74 – 1.85 |
| Random: susceptible biased-medium | 1.91 | 1.85 – 1.97 |
| **Random Effects** |  |  |
| $\sigma^2$ | 1.77 | |
| $\tau_{00}$ number_infections:(fox_uniqueID:run) | 1.40 | |
| $\tau_{00}$ fox_uniqueID:run | 0.01 | |
| $\tau_{00}$ run | 0.00 | |
| $N_{\text{number\_infections}}$ | 19 | |
| $N_{\text{fox\_uniqueID}}$ | 17424 | |
| $N_{\text{run}}$ | 1089 | |
| Observations | 122609 |  |

The landcover-based movement behaviour resulted in a greater number of effective contacts compared to the random movement behaviour. Within this landcover-based movement scenario, the greatest number of effective contacts was observed when there was an equal probability of susceptible and infected foxes leaving a park at a low and medium intensity, 754.17 (±148.85) contacts and 763.39 (±110.86) contacts respectively. In addition, the proportion of effective contact events that occurred outside of the parks was greater than the proportion of contact events that occurred within the two parks, as 67%-78% of contacts occurred along movement routes (Figure 3(b); Supplementary Material: Table S1). The model for the number of effective contact events had an *R^2^* = 0.96 (Table 4). Movement behaviour and the probability of leaving a park according to infection status had a significant effect on the number of effective contact events (Supplementary Material: Table S3).

**Table 4.**
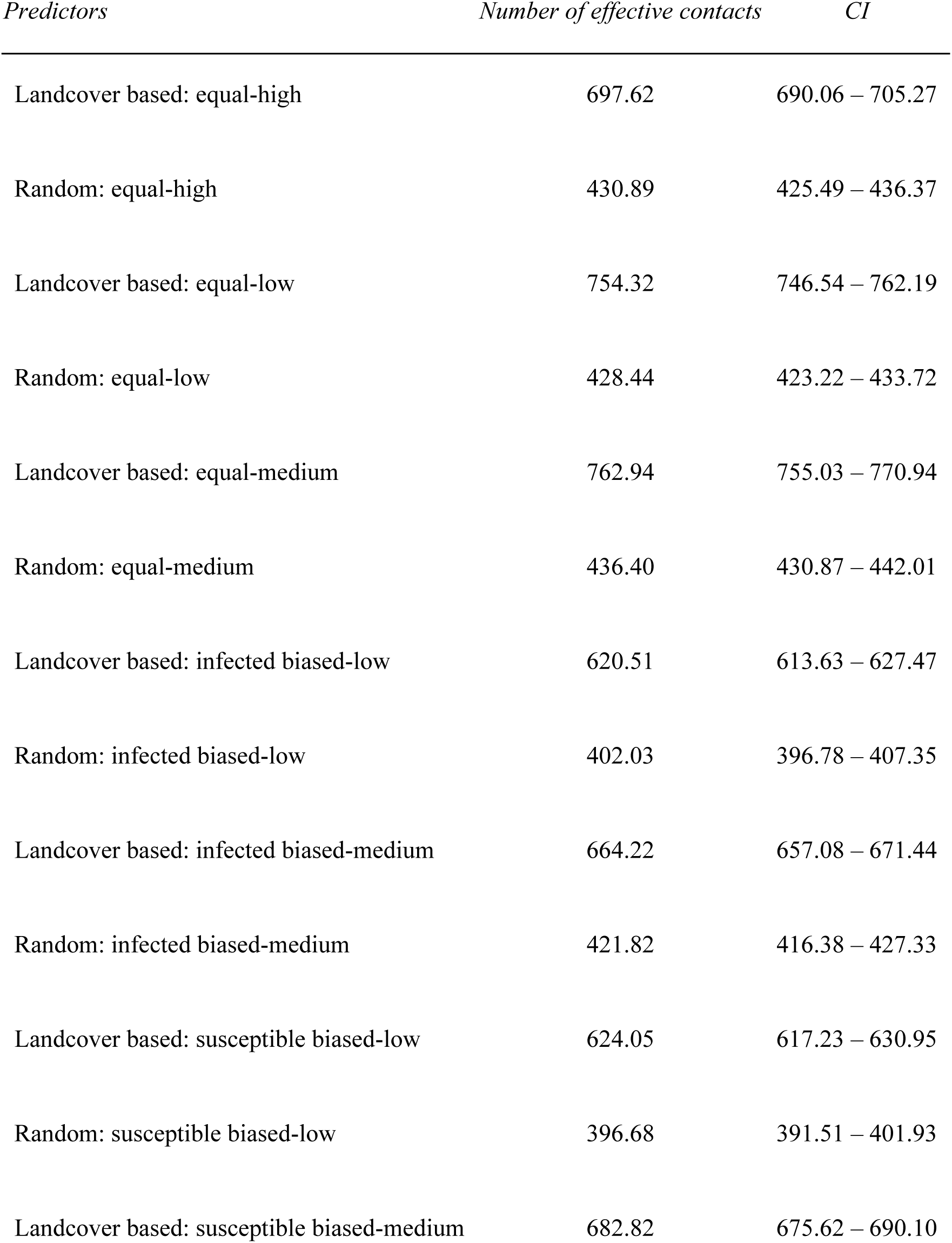
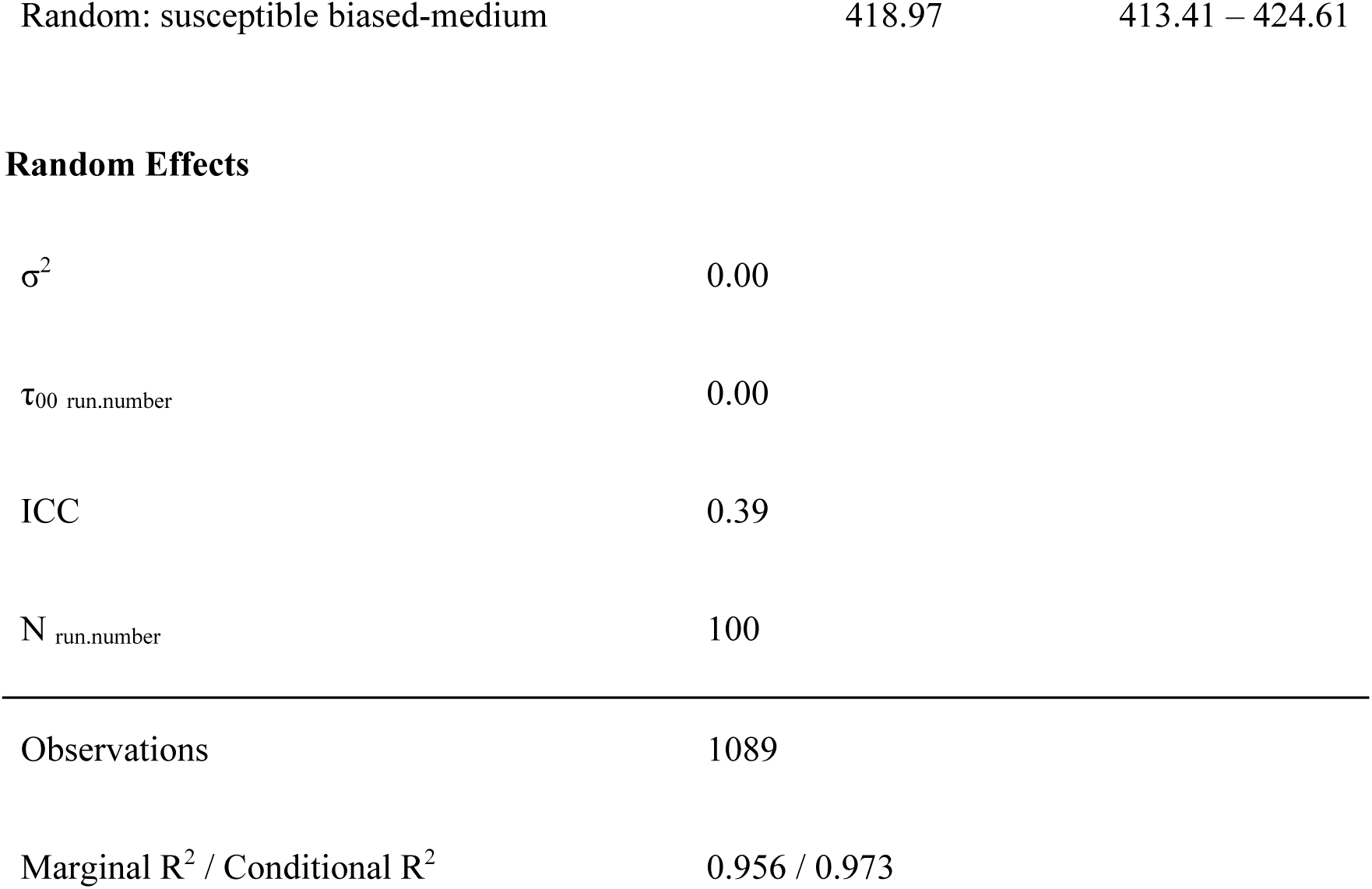
Results for the number of effective contact events from the general mixed model.

| *Predictors* | *Number of effective contacts* | *CI* |
| --- | --- | --- |
| Landcover based: equal-high | 697.62 | 690.06 – 705.27 |
| Random: equal-high | 430.89 | 425.49 – 436.37 |
| Landcover based: equal-low | 754.32 | 746.54 – 762.19 |
| Random: equal-low | 428.44 | 423.22 – 433.72 |
| Landcover based: equal-medium | 762.94 | 755.03 – 770.94 |
| Random: equal-medium | 436.40 | 430.87 – 442.01 |
| Landcover based: infected biased-low | 620.51 | 613.63 – 627.47 |
| Random: infected biased-low | 402.03 | 396.78 – 407.35 |
| Landcover based: infected biased-medium | 664.22 | 657.08 – 671.44 |
| Random: infected biased-medium | 421.82 | 416.38 – 427.33 |
| Landcover based: susceptible biased-low | 624.05 | 617.23 – 630.95 |
| Random: susceptible biased-low | 396.68 | 391.51 – 401.93 |
| Landcover based: susceptible biased-medium | 682.82 | 675.62 – 690.10 |
| Random: susceptible biased-medium | 418.97 | 413.41 – 424.61 |

**Random Effects**
|  |  |
| --- | --- |
| $\sigma^2$ | 0.00 |
| $\tau_{00}$ run.number | 0.00 |
| ICC | 0.39 |
| $N$ run.number | 100 |

|  |  |
| --- | --- |
| Observations | 1089 |
| Marginal $R^2$ / Conditional $R^2$ | 0.956 / 0.973 |

Overall, we found that both the probability of leaving a park and landcover-based movement behaviour had a statistically significant effect on the *R_e_* and the number of effective contacts in our study area. These values were greatest when foxes utilized the landcover-based movement behaviour and when susceptible and infected foxes had an equal probability of leaving their initial park.

## 4 Discussion

The results obtained with our MTM model provide evidence that both movement behaviour informed by landcover type and movement ability of foxes, affect the *R_e_* and the number of effective contacts. In particular, movement along least-cost paths and an equal probability of leaving a park of susceptible and infected foxes lead to increased encounters and subsequent spread of mange.

The MTM that included the landcover-based movement behaviour led to a greater mean *R_e_*, and an increase in effective contact rates between susceptible and infected individuals. These results suggest that the dynamics of mange may be altered by the landscape, and can be exacerbated in urban areas, where fox movement is highly dependent on the degree of landcover permeability to movement (Lovell et al. 2022). Foxes may be more prone to encountering mange-infected individuals along urban corridors that facilitate movement between fragmented habitat patches. These findings are supported by observational studies of mange infected foxes and coyotes, which were more likely to utilize roads and anthropogenic resources (Reddell et al. 2023; Wails et al. 2024). In addition, when looking at the proportion of contact events observed, the percentage of effective contact events in our study area was greater when foxes were moving between parks than within, as the individuals tended to be funneled into a single movement corridor within the study area. This suggests that movement behaviour across the matrix, hence the ability to move through different land cover types and their spatial configuration, plays a significant role in the transmission of mange in heterogenous landscapes. For example, the presence of more or less movement corridors within a study area will likely influence contact rates and disease transmission. These results are in accordance with previous studies which have found that disease is more likely to be sustained in models where movement decisions are mechanistically considered (Scherer et al. 2020).

Empirical studies assessing the prevalence of mange in urban foxes have noted that infection occurs in clusters and can be affected by human presence and the level of green spaces in the city (Newman et al. 2002; Carricondo-Sanchez et al. 2017). The results observed here, namely increased transmission within movement corridors, may support the notion that increased frequency of contact between hosts, namely by the use of common movement corridors could facilitate mange spread. This is in support of Hess’s metapopulation models (Hess 1996) which suggest an increased risk of disease spread particularly along movement corridors within home ranges, as in the model depicted here. Our sample size of 16 foxes within a 1km^2^ study area was higher than what has been observed on average, which may have further led to higher transmission rates (Scott et al. 2014), however, the total number of fox individuals remains constant across scenarios.

The greatest mean *R_e_* and effective contact events were observed when susceptible and infected foxes had an equal probability of leaving a park and when movement was landcover-based. This suggests that when there is equal mixing between susceptible and infected foxes regardless of animal density within habitat areas, mange is likely to spread at a higher rate following an increase in contact rates as individuals move around the city. This supports previous findings suggesting mange transmission is predominantly frequency-dependent (Devenish-Nelson et al. 2014). In further support of this, we expected that the susceptible-biased simulations, where infected individuals were less likely to exit green areas, would lead to lower *R_e_* and effective contact events than in the infected-biased scenario, as previous work has shown that reduced mobility of hosts may lead to less sustained transmission (Ichinose et al. 2018).Instead, our results suggest that limited habitat availability in cities, leading to increased overlap in space use and hence frequency of encounter rates, can lead to higher disease transmission rates, regardless of whether infected foxes tend to remain in such patches. Future studies should explicitly examine the effects of fox density and different habitat availabilities in urban landscapes on the spread of mange.

The MTM purposely omitted demographic factors of foxes, such as births and deaths, as well as disease-induced mortality. This was done to directly test the effects of movement behaviours and ability on the metrics of mange spread, avoiding additional complexity in the analysis of the results (Tracey et al. 2014). The inclusion of these demographic factors may lead to a more accurate estimation of the *R_e_* given that individuals are likely to be added and removed from the population at varying rates, affecting the number of effective contact events. While there is little evidence of immunity to mange following infection, some treatments that can provide temporary immunity to fox populations have been studied (Foley et al. 2023; Rudd et al. 2020). If successful, such treatments can potentially lead to an additional compartment of recovered individuals, and further impact mange dynamics. In addition, including behavioural variables in movement decisions according to age and sex structure may also improve the model. Previous studies have observed differential probability of mange transmission according to age, with juveniles being more likely to contract and spread the disease (Devenish-Nelson et al. 2014; Pisano et al. 2019; Scott et al. 2020). It is therefore possible that juveniles could have different movement patterns or interaction behaviours than adults, influencing the probability of them coming into direct contact with other individuals. Further work is necessary to differentiate the movement patterns of juveniles from adults, as their inclusion in models of mange could be important for control efforts. Understanding the social dynamics and structure of foxes on a temporal scale could also be useful to better quantify the dynamics of mange (Albery et al. 2021). Temporal dynamics may also be important to consider as there is evidence that interactions between foxes from different social groups may vary based on sex and season (Dorning & Harris 2019). Contact rates between foxes tend to increase during the breeding season in the summer and decrease during the winter (Dorning & Harris 2019). These behaviours may associate with differential levels of mange transmission on a temporal scale.

Our model only considered intraspecific mange transmission, however, it is possible that mange dynamics could be exacerbated in fox populations through interspecific transmission from other mammals (Cypher et al. 2017). Foxes are more likely to have habitat overlaps in cities with other species that are prone to mange infection such as raccoons and coyotes (Rentería-Solís et al. 2014; Mueller et al. 2018; Peterson et al. 2021). It is unclear whether transmission between these animals to foxes occurs, as studies suggest the *S. scabiei* mite may be host-specific (Arlian & Morgan 2017). Yet, mange transmission events through predator-prey interactions have been observed in wild animal populations in Africa (Gakuya et al. 2011). In addition, there is evidence of indirect fomite transmission between different carnivore species in denning sites (Kołodziej-Sobocińska et al. 2014). Therefore, it is important to consider alternate routes of transmission that could result from interactions with other host species. In addition, indirect transmission may also occur, in particular through the shedding of mites in dens and subsequent den-sharing, which could further amplify transmission (Loredo et al. 2020). Future efforts should focus on understanding the host-specificity of the *S. scabiei* mite and the role den-sharing plays in transmission, which could lead to a reduction in urban fox population sizes.

Despite these limitations, this study stresses the importance of including realistic movement behaviours in the spread of mange. Unlike previous models of mange (Lunelli 2010; Devenish-Nelson et al. 2014; Alexander et al. 2016), the MTM is novel in that it includes spatially explicit properties of transmission based on knowledge-based assumptions of mange spread and movement of foxes following a landcover-based resistance matrix. This study highlights the significance of landscape configuration, movement behaviour, and movement ability of foxes in sustaining the transmission of mange in urban settings. Future directions that could improve model results would be model validation through fox movement tracking and collection of epidemiological data on mange infections, as well as extending the model to a larger spatial scale (Walton et al. 2018).

Nonetheless, our results emphasize the role of movement corridors in disease transmission through direct contact, especially in fragmented landscapes. However, our results are also applicable to disease transmission risk in movement corridors between fragmented forested landscape outside of urban areas, as previous research shows increased movement within movement corridors across most species (Pither, 2023). Furthermore, our results may be also applicable to vector-borne diseases where low-mobile vectors are shed, and successful transmission depends on shared space use of hosts. For example, a higher density of ticks has been found along movement corridors, within and outside urban areas (Lilly et al. 2025).

As urbanization continues to accelerate, habitats will become further fragmented (Haddad et al. 2015), which our findings suggest will lead to increased encounter rates with infected animal hosts. Therefore, understanding the movement routes of wildlife species that contribute to the spread of disease, and the alteration of population and community dynamics will become increasingly important. ABMs like the one presented here are powerful tools to better quantify the dynamics of direct-contact diseases when doing so empirically is challenging and can better inform disease control and conservation efforts (Watkins et al. 2011; Miller et al. 2014; Belsare & Stewart 2020).

## 5 Conclusions

This study builds on previous modelling efforts of the spread of mange in urban settings, while integrating spatially explicit movement behaviours according to landcover types and infection status. Our results show that movement of foxes following paths of least resistance through a heterogeneous landscape and non-biased movement behaviour of susceptible and infected individuals leads to a greater number of effective contact events and *R_e_*in comparison to an MTM with random movement. Unexpectedly, the infected-biased and susceptible-biased scenarios resulted in a similar number of contact events and *R_e_*, likely due to increased infection events across movement corridors. Therefore, our results highlight that fragmentation may lead to an increase in transmission rates following the canalized movement of hosts along corridors, and that disease-induced differential likelihood of leaving parks may alter disease dynamics, reducing transmission when movement abilities between susceptible and infected hosts are imbalanced.

## Declarations

## Ethical Approval

Not applicable

## Authors’ Contributions

N.D. wrote the main manuscript text. M.-J.F., T.G.C., and M.K. contributed edits throughout the writing process. N.D. prepared the figures and tables. T.G.C. and M.K. contributed to writing data cleaning and analysis scripts. All authors reviewed the manuscript.

## Availability of data and materials

All data and code are available on Figshare at: 10.6084/m9.figshare.24501934.

## Acknowledgements

We would like to thank Drs. Nicole Mideo, Donald C. Jackson, and Benjamin Gilbert for their guidance on the study design and concepts. Thanks to Dr Paul Smaldino for providing help with the NetLogo movement procedure code.

## 6 Appendix

### 6.1 ODD Protocol

#### Overview

##### 6.1.1 Model Purpose

The MTM is used to quantify sarcoptic mange effective contact events and *R_e_* in the urban region of Scarborough, according to two types of between-park movement behaviours: random and landcover-based. Moreover, the MTM tests how different probabilities of leaving a park according to infection status will influence mange transmission. The model includes two low-level entities: individual adult foxes and habitat cells described by landcover types and resistance to movement values. Movement behaviours occur based on a decision-making process according to the location of a fox on the landscape (Supplementary Material: Fig 4).

##### 6.1.2 State Variables

Low-level entities: Individual foxes described as adults (no sex or age structure) and geographical cells described according to abiotic factors (landcover type, resistance to movement).

High-level entities: A population of 16 foxes.

##### 6.1.3 Scales

Size of habitat cells: 10×10m.

Length of time steps: 100 time steps/day.

Simulation duration: 150000 time-steps (1500 days).

##### 6.1.4 Process Overview

Between-park fox movement: according to two types of movement: random and landcover-based. Within-park fox movement: according to a correlated random walk (CRW) (Towertown et al. 2016).

Mange spread: according to the probability of infection by a contact.

##### 6.1.5 Scheduling

Time handling: Time is discrete with movements being updated daily with 100 time steps per day. 100 time steps are chosen as the cell size in the model is 10×10m, and daily movement distances of red foxes within cities are estimated to be around 1km (Rosatte & Allen 2009).

Model processes groupings: Movement and disease spread begin in a synchronous fashion from the initialization of the model.

#### Design Concepts

Expectation: Individuals will choose cells with the lowest resistance values for between-park movement as determined by landcover type (LaRue & Nielsen 2008).

Emergence: Number of effective contact events as determined by the movement of infected and susceptible host species and the effective reproduction number *R_e_*.

Sensing: Individuals perceive the resistance costs of all eight neighbouring cells and move to the lowest value while maintaining a global direction.

Stochasticity: Initial disease state of foxes; location of foxes within the park at the start of the simulation; probability of becoming infected; latency period; infectious period; probability of moving out of a park.

#### Details

##### 6.1.6 Initialization

The simulation starts with a set number of foxes (*n*=16), which represents a realistic density of foxes per km^2^ in cities, with one infected fox at the (I – infected stage) beginning of the simulation and all other foxes at the susceptible stage (S) (Marks & Bloomfield 1999). Foxes begin in two parks with an equal number of individuals (8 per park). Each fox moves within the park using a CRW. Foxes who hit a park edge then move to the next park using random movement or landcover-based movement by selecting cells with the lowest values of resistance.

##### 6.1.7 Submodels

###### Movement within Parks

Foxes move within a park using a correlated random walk sampled from a Weibull distribution (Morales et al. 2004). The turning angle at each given step (step size: 1 cell) is sampled from a cosine function, with a mean turning angle of 180 degrees, such that a forward direction is maintained. Correlation in movement is high with a probability of 0.9 of retaining the same direction at the next timestep (Supplementary Material: Fig S1).

###### Movement Between Parks

Movement between parks occurs according to one of two behaviours, random movement, and landcover-based movement. In the random movement behaviour, if foxes hit a park edge, they will move out of the park at a given probability of leaving a park according to infection status. Foxes will maintain a global direction towards the next park and will move out of a park by selecting a random cell at each time step.

In the landcover-based movement model, foxes will move out of a park at a given probability of leaving a park according to infection status. Foxes will maintain a global direction towards the next park and will perceive of all eight neighbouring cells. They will use the cells with the lowest resistance to get to their destination (i.e., the next park). If the lowest resistance value cell is in a direction opposite to the global direction, they will favour the cell that maintains the global direction over the cell with the lowest resistance value (Kanagaraj et al. 2013).

###### Mange Transmission

Susceptible foxes acquire the disease at a given transmission probability if an infected fox is within a radius of 1 cell of their location. They will then move onto the exposed stage (E) and will become infected (I) after an exponentially distributed exposure period. Foxes will recover and join the susceptible pool (S) again after an exponentially distributed infectious period (Supplementary Material: Fig S2). Mode of transmission is not explicitly incorporated in the model, with it expected to arise as an emergent property.

##### 6.1.8 Submodel Parameters

###### Movement Within Park Submodel

Foxes within a park move according to a correlated random walk (CRW). The correlated random walk is sampled from a Weibull distribution, with the angle of movement sampled from a cosine function (Jonhson et al. 2008). Because foxes maintain a high correlation in movement, the correlation in movement probability for the next step is set at 0.9 (Towerton et al. 2016). To keep a forward direction, the mean turn angle is set to 180 degrees.

###### Movement Between Parks Submodel

Foxes between parks move randomly, or according to the resistance values summarized in Table 1.

###### Mange Transmission Submodel

Mange in red foxes has a main transmission route of direct contact, with indirect transmission through fomites being considered negligible (Carricondo-Sanchez et al. 2017). For this reason, only direct contact transmission is considered in the model. Foxes that are susceptible and found within a radius of 1 cell of an infected fox have a probability of getting infected. Assessing data from previous mange outbreaks reveals a high attack rate at the beginning of the outbreak, ranging from 45% - 70% (Carricondo-Sanchez et al. 2017; Pisano et al. 2019; Scott et al. 2020). This attack rate was used in order to assess an appropriate infection probability for the simulation duration, resulting in an infection probability of about 0.03 per day for 1500 days or 150000 time steps.

Once a fox becomes infected, it enters an exposure period during which it is asymptomatic and not infectious. The exposure and infectious periods are exponentially distributed with an average of 30 and 120 days, respectively (Devenish-Nelson et al. 2014).

