## Supplementary Material for "The effects of fox movement and landscape heterogeneity on the spread of sarcoptic mange in urban settings"

**Table S1** – Summary results for the Scarborough region MTM (10×10m cell resolution): The number of simulations where the spread is sustained through the simulation time, the mean $R_e$, the mean # of effective contacts, mean within-park # of contacts (%), and mean between-park # of contacts (%) summarized for the simulations where the spread is sustained. Results in a regular font are for the random movement behaviour, while italicized results are for the landcover-based movement behaviour.

| Probability of leaving a park | Sustained spread (/100) | Mean $R_e$ (Standard deviation) | Mean # of effective contacts (Standard deviation) | Mean within-park # of contacts (%) | Mean between-park # of contacts (%) |
|---|---|---|---|---|---|
| Equal: Low | 85 | 2.21 (0.35) | 428.06 (29.36) | 45.53 | 54.47 |
| | *81* | *3.7 (0.86)* | *754.17 (148.85)* | *33.35* | *66.65* |
| Equal: Medium | 75 | 2.19 (0.4) | 435.65 (28.19) | 33.45 | 66.55 |
| | *79* | *3.85 (0.7)* | *763.39 (110.86)* | *23.9* | *76.1* |
| Equal: High | 78 | 2.13 (0.24) | 430.97 (31.41) | 30.21 | 69.79 |
| | *75* | *3.47 (0.57)* | *696.99 (74.74)* | *21.75* | *78.25* |
| Infected-biased: Low | 75 | 2.01 (0.32) | 401.51 (25.44) | 35.33 | 64.67 |
| | *78* | *3.11 (0.52)* | *621.38 (67.87)* | *27.04* | *72.96* |
| Infected-biased: Medium | 74 | 2.15 (0.29) | 422.24 (25.58) | 31.06 | 68.94 |
| | *80* | *3.39 (0.61)* | *666.73 (77.83)* | *23.81* | *76.19* |
| Susceptible-biased: Low | 76 | 2.04 (0.32) | 396.58 (28.94) | 35.56 | 64.44 |
| | *81* | *3.13 (0.58)* | *623.62 (87.14)* | *27.46* | *72.54* |
| Susceptible-biased: Medium | 69 | 2.09 (0.33) | 419.36 (32.13) | 30.89 | 69.11 |
| | *83* | *3.3 (0.43)* | *682.37 (71.3)* | *23.2* | *76.8* |

**Table S2** – Analysis of Deviance results for the $R_e$ general mixed model. The treatment term represents a combination of the probability of leaving patch scenarios and the between patch movement behaviour: (1) random or (2) landcover-based. Significant values are denoted with an asterisk (***).

| Term | df | chi-squared statistic | P-value |
|---|---|---|---|
| Treatment | 14 | 44621 | **< 2.2e-16*** ** |

**Table S3** – Analysis of Deviance results for the number of effective contact events linear mixed model. The scenario term represents the probability of leaving a patch scenario, the movement behaviour term represents the between patch movement behaviour: (1) random or (2) landcover-based, and the last term represents an interaction of the two. Significant values are denoted with an asterisk (***).

| Term | df | Chi-Square Statistic | *p*-value |
|---|---|---|---|
| Treatment | 14 | 3149201 | < 2.2e-16*** |

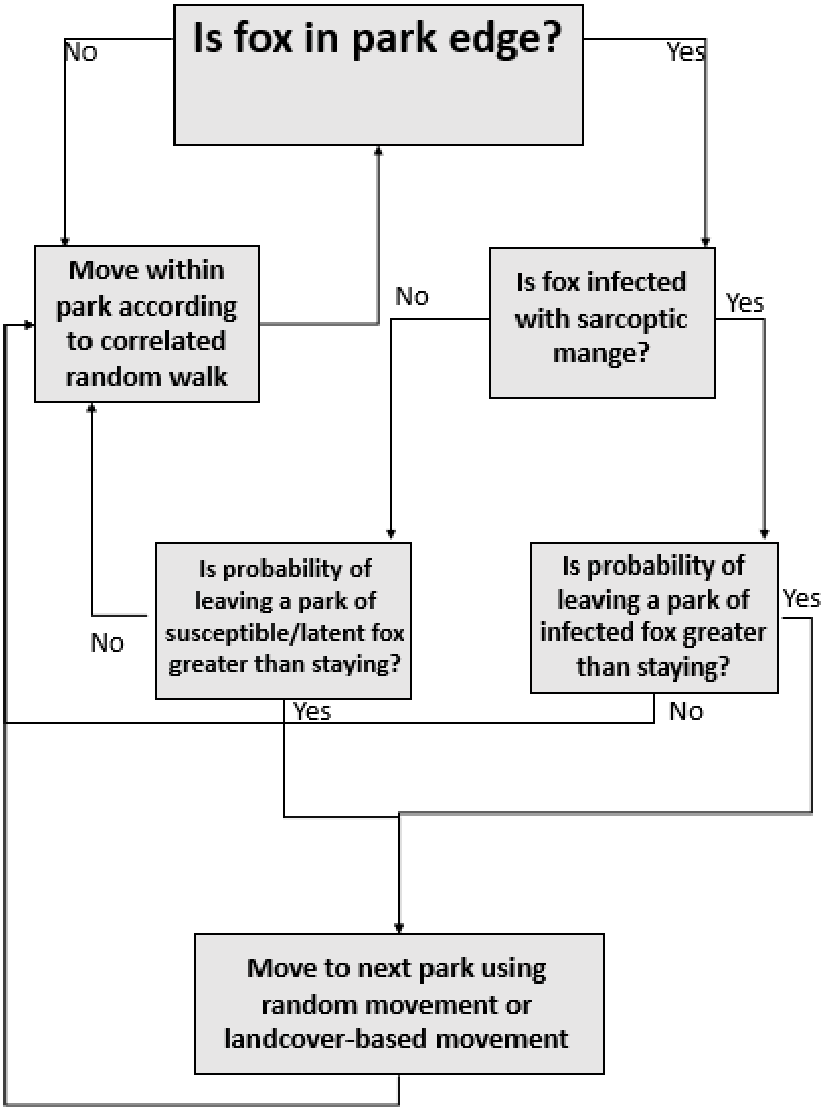


**Figure S1** – Flow chart diagram of the between-park movement and within-park movement procedures of foxes.

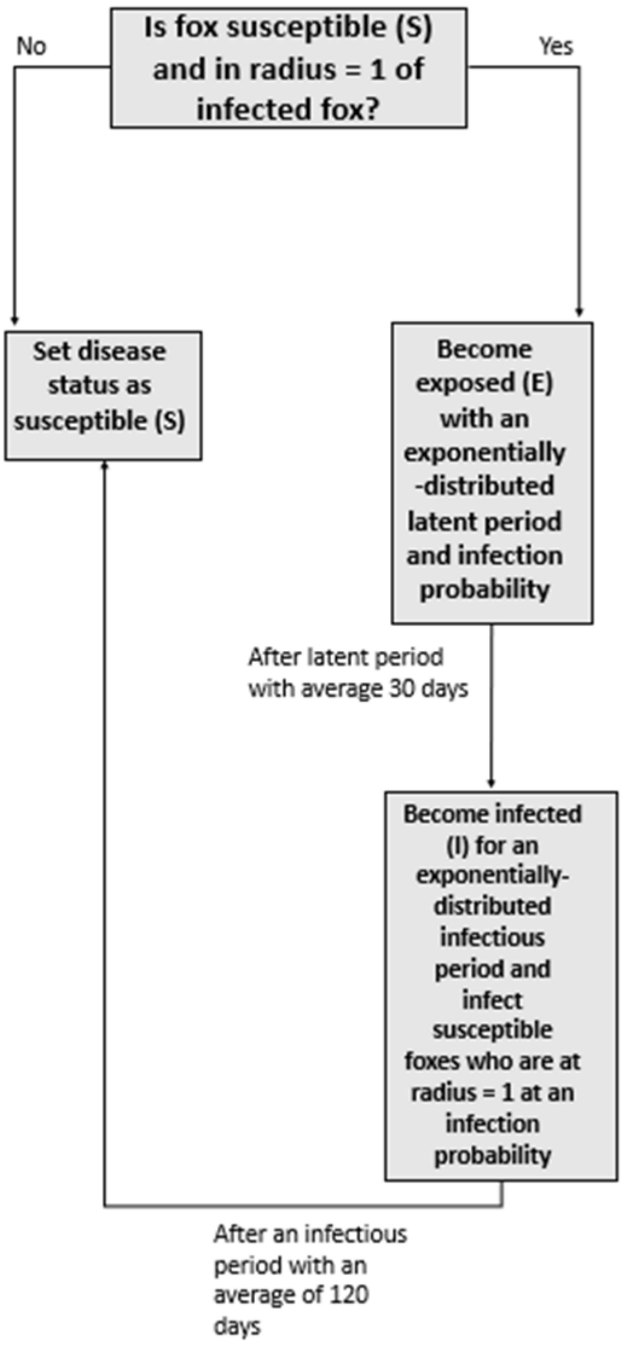


**Figure S2** – Flow chart diagram of the sarcoptic mange SEI disease procedure.
